# The human HELLS chromatin remodelling protein promotes end resection to facilitate homologous recombination within heterochromatin

**DOI:** 10.1101/504043

**Authors:** G. Kollarovic, C. E. Topping, E. P. Shaw, A. L. Chambers

## Abstract

Efficient double-strand break repair in eukaryotes requires manipulation of chromatin structure. ATP-dependent chromatin remodelling enzymes can facilitate different DNA repair pathways, during different stages of the cell cycle and in a range of chromatin environments. The contribution of remodelling factors to break repair within heterochromatin during G2 is unclear.

The human HELLS protein is a Snf2-like chromatin remodeller family member and is mutated or misregulated in several cancers and some cases of ICF syndrome. HELLS has been implicated in the DNA damage response, but its mechanistic function in repair is not well understood. We find that HELLS facilitates homologous recombination at two-ended breaks within heterochromatic regions during G2. HELLS enables end-resection and accumulation of CtIP at IR-induced foci. We identify an interaction between HELLS and CtIP and establish that the ATPase domain of HELLS is required to promote DSB repair. This function of HELLS in maintenance of genome stability is likely to contribute to its role in cancer biology and demonstrates that different chromatin remodelling activities are required for efficient repair in specific genomic contexts.

## INTRODUCTION

Unrepaired double-strand breaks (DSBs) can lead to cytotoxicity, whilst misrepair of breaks can result in genomic instability that may promote tumourigenesis (Jackson and Bartek, 2009). There are two major pathways of DSB repair (DSBR)-canonical non-homologous end joining (c-NHEJ) and homologous recombination (HR). HR requires a sister chromatid to provide a template for repair and therefore is restricted to the S and G2 phases of the cell cycle (Filippo *et al*., 2008). HR occurs with slower kinetics than NHEJ, is error-free and performs an essential function in repair of one-ended breaks generated by collapse of replication forks during S-phase, as well as contributing to repair of two-ended breaks in G2 (Trenz *et al*., 2006; Beucher *et al*., 2009). HR begins with break recognition and initiation of end-resection by the MRE11-RAD50-NBS1 (MRN) complex in association with CtIP (Limbo *et al*., 2007; Sartori *et al*., 2007; You *et al*., 2009). Further extensive resection by EXO1 or BLM/DNA2 (Gravel *et al*., 2008) results in 3’ single-stranded DNA (ssDNA) overhangs that are initially coated in RPA (Replication Protein A). BRCA2 promotes replacement of RPA with RAD51, to produce a RAD51 nucleofilament that mediates homology searching and strand invasion.

In eukaryotes, DSBR occurs in a chromatin environment, which can act as a barrier to repair. Transcriptionally repressed and repetitive regions of the genome are packaged into compact, condensed heterochromatin (Allshire and Madhani, 2018). Repair of DSBs within heterochromatin occurs more slowly than breaks located in euchromatin during both G0/G1 and G2 (Goodarzi *et al*., 2008; Woodbine *et al*., 2011) and the mutation rate is higher within heterochromatin, in part due to its repetitive nature (Schuster-Böckler *et al*., 2012). In G1 heterochromatic breaks are repaired by NHEJ and in G2 by HR, (Riballo *et al*., 2004; Beucher *et al*., 2009; Lobrich *et al*., 2010). In both instances heterochromatin repair correlates with a slow component of repair (∼15% of DSBs). The DNA damage response kinase, ATM has been shown to play a role in overcoming the barrier to repair posed by heterochromatin in both G0/G1 cells and G2 (Goodarzi *et al*., 2008; Beucher *et al*., 2009).

ATP-dependent chromatin remodelling enzymes, couple ATP hydrolysis to translocation along DNA to alter nucleosome composition or position (Clapier *et al*., 2017). Chromatin remodelling is necessary for efficient DSBR (Jeggo and Downs, 2014). INO80, SWR1, ISW1 and SWI/SNF complexes are recruited to DSBs and promote repair, whilst CHD1, SRCAP and SMARCAD1 have been associated with end-resection (reviewed in Jeggo and Downs, 2014). Questions remain over whether each remodeller is required at every break to enable different stages of repair, or if specific remodellers facilitate repair of subsets of breaks, for instance those with greater damage complexity or within different chromatin contexts. During G0/G1, efficient repair within heterochromatin requires dispersal of CHD3 (Goodarzi, Kurka and Jeggo, 2011) and activity of the ACF1-SNF2H/SMARCA5 chromatin remodelling complex (Klement *et al*., 2014). However, involvement of ATP-dependent chromatin remodelling in repair of heterochromatic breaks during G2, is poorly understood.

Sequence analysis identifies the human HELLS protein (also known as LSH, PASG or SMARCA6) as a Snf2-like chromatin remodelling enzyme (Geiman, Durum and Muegge, 1998; Lee *et al*., 2000; Raabe *et al*., 2001; Flaus *et al*., 2006). The ATPase of HELLS is stimulated by nucleosomes and remodelling activity has been demonstrated for a HELLS-CDC7a complex, with the *Drosophila* homolog, Ddm1, also shown to slide nucleosomes *in vitro* (Brzeski and Jerzmanowski, 2003; Burrage *et al*., 2012; Jenness *et al*., 2018). HELLS is ubiquitously expressed, with highest levels found in proliferating tissues active in recombination such as the thymus and testes (Geiman, Durum and Muegge, 1998; Lee *et al*., 2000; Raabe *et al*., 2001). HELLS is mutated or misregulated in tumour samples from several cancer types (Yano *et al*., 2004; Kim *et al*., 2010; Janus *et al*., 2011; Keyes *et al*., 2011; Von Eyss *et al*., 2012) and is also mutated in some cases of immunodeficiency-centromeric instability-facial anomalies syndrome (ICF), characterised by instability within pericentromeric repeats (Thijssen *et al*., 2015). HELLS-deficient mice either age prematurely or die shortly after birth, while HELLS mutant MEFs (mouse embryonic fibroblasts) display premature onset of senescence, reduced proliferation, aberrant chromosome segregation and increased DNA content (Fan *et al*., 2003; Sun *et al*., 2004; Von Eyss *et al*., 2012). HELLS-deficient MEFs have reduced global DNA methylation and it has been proposed that HELLS permits recruitment of DNA methyltransferases for *de novo* DNA methylation, particularly at repetitive sequences (Dennis *et al*., 2001; Yan *et al*., 2003; Muegge, 2005; Myant and Stancheva, 2008; Von Eyss *et al*., 2012). Budding yeast don’t methylate their DNA, but possess a HELLS homolog (Irc5), raising the possibility of another conserved function for this subfamily of chromatin remodelling enzymes (Alvaro, Lisby and Rothstein, 2007; Litwin *et al*., 2017). Indeed, a role for HELLS in maintenance of genome stability has emerged. HELLS-deficient MEFs and MRC5 human fibroblasts display sensitivity to DNA damaging agents, inefficient repair and aberrant DNA damage responses (Burrage *et al*., 2012). However, the mechanistic role of HELLS in protection of genome integrity is not yet fully described. Two unanswered key questions are: in which contexts or genomic regions does HELLS contribute to repair, and which step of DNA repair is impacted by HELLS?

In this study, we find that the human HELLS protein promotes genome stability following IR and as part of the response to spontaneously occurring damage. We find that HELLS facilitates DSBR by HR during G2 and is required for efficient repair of breaks within heterochromatin. Our data show that HELLS impacts DNA end-resection and promotes accumulation of CtIP in IR-induced foci. We uncover an interaction between HELLS and CtIP and establish that HELLS ATPase activity contributes to its function in DSBR. Our data reveal a role for the HELLS chromatin remodeller in repair of heterochromatic DSBs in G2 and provide insights into the function of HELLS in preventing genome instability.

## RESULTS

### HELLS promotes genome stability

To investigate the function of the human HELLS protein in maintenance of genome stability, we established conditions to deplete HELLS using siRNA in hTert-immortalised normal human fibroblasts (1BR-hTert) (Figure 1A). Phosphorylated histone H2AX foci (γ-H2AX), a marker of DSBs, were visualised in G2 cells identified by their pan-nuclear CENP-F staining (Kao, McKenna and Yen, 2001) (Figure 1B). Unexpectedly, HELLS depletion resulted in elevated numbers of γ-H2AX foci in undamaged, asynchronously growing G2 cells (Figure 1C and Figure S1A). An increase in spontaneous γ-H2AX foci did not occur in serum-starved cells downregulated for HELLS (Figure S1B). HELLS-depleted fibroblasts did not show significant changes in cell cycle profile and retained functional G1/S and G2/M checkpoints (Figure S1C). The greater number of γ-H2AX foci is consistent with either increased spontaneous break formation, or a defect in repair of DSBs on HELLS downregulation.

**Figure 1.**
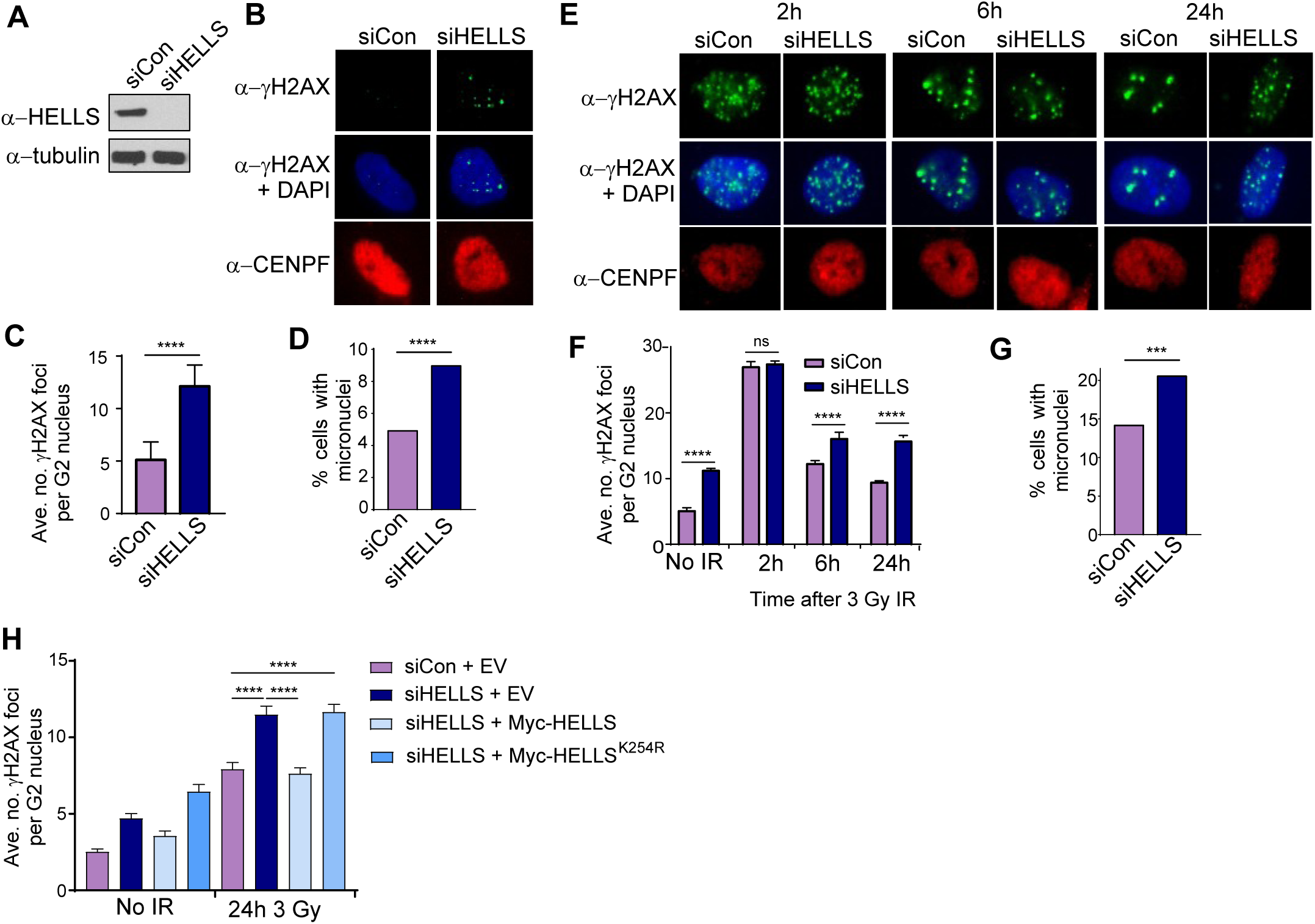
HELLS promotes genome stability in undamaged cells and following IR. **(A)** Western blot analysis of efficiency of HELLS depletion by siRNA in 1BR-hTert cells. Anti-tubulin was used as a loading control. **(B)** Representative immunofluorescent images of spontaneously arising γ-H2AX foci in 1BR-hTert cells transfected with the indicated siRNA. CENPF staining was used to identify G2 cells. **(C)** Quantification of spontaneous γ-H2AX foci in G2 nuclei of 1BR-hTert cells transfected with indicated siRNA. Data are from 6 independent experiments. **(D)** Quantification of micronuclei in undamaged 1BR-hTert cells transfected with indicated siRNA. A minimum of 1100 nuclei were analysed, from three independent experiments. **(E)** Representative immunofluorescent images treated with the indicated siRNA at different time points after exposure to 3 Gy IR. **(F)** Quantification of clearance of γ-H2AX foci following 3 Gy irradiation in 1BR-hTert cells treated with indicated siRNA. Data from three independent experiments. **(G)** Quantification of micronuclei in 1BR-hTert cells transfected with indicated siRNA after 6 Gy IR. A minimum of 715 nuclei were analysed, from three independent experiments. **(H)** siRNA-resistant WT or K254R mutant constructs were introduced to siHELLS cells and γ-H2AX foci numbers quantified 24 h following 3 Gy IR. Data from a minimum of three independent experiments. Data shown in panels C, F and H are mean +/- sem. **** indicates a p-value of <0.0001, and ns indicates not significant by unpaired two-tailed t-test analysis.

The incidence of spontaneous micronuclei was also elevated in cells downregulated for HELLS (Figure 1D). Micronuclei result when a chromosome or chromosome fragment is not incorporated into the daughter nuclei during mitosis and can arise due to chromosome segregation defects or the presence of unrepaired or misrepaired breaks. Our data from HELLS-depleted fibroblasts are consistent with observations in LSH -/- MEFs (the murine homologue of HELLS) which showed elevated micronuclei formation and chromosomal instability (Fan *et al*., 2003). These data support a function for the human HELLS protein in protection of genome integrity.

### HELLS depletion results in genome instability following exposure to IR

To examine the role of HELLS in DSBR, cells were irradiated with 3 Gy IR and numbers of γ-H2AX foci in G2 cells analysed over time (Figure 1E). Addition of aphidicolin at the time of irradiation precluded cells damaged during G1 progressing into G2 and prevented one-ended breaks arising during S-phase from entering our analysis (Beucher *et al*., 2009; Lobrich *et al*., 2010). Aphidicolin has been shown to not affect NHEJ, HR or numbers of γ-H2AX foci (Beucher *et al*., 2009; Shibata *et al*., 2011) and the cell cycle profile of HELLS-depleted cells 24 hours after aphidicolin, irradiation or both, was the same as control cells (Figure S1D). S-phase cells were excluded based on their pan-nuclear, aphidicolin-dependent H2AX phosphorylation and CENPF-positive staining. Consequently, we are examining the effect of HELLS on repair of two-ended DSBs that are generated and repaired during G2. HELLS downregulated cells possessed comparable numbers of γ-H2AX foci to cells treated with control siRNA 2 hours after irradiation, but at later times after IR (6 and 24h) HELLS-depleted cells displayed increased numbers of persistent γ-H2AX foci (Figures 1F and S1E). These data suggest that break formation by IR and rapid DSBR is independent of HELLS, but that HELLS depletion results in a defect in the slower component of repair in G2 cells. This phenotype is reminiscent of the late DSBR in cells deficient in ATM, Artemis, RAD51 or BRCA2 (Beucher *et al*., 2009). HELLS localises to the nuclear fraction and levels did not increase following IR (Figure S1F). Consistent with a function for HELLS in repair of DSBs, irradiated HELLS-deficient cells also exhibited increased frequency of micronuclei compared with control cells (Figures 1G). Together these data suggest a role for the human HELLS protein in promoting timely DSB repair and maintenance of genome stability.

### The ATP-binding site of HELLS is necessary for its function in DSBR in G2

To confirm the role of HELLS in γ-H2AX clearance and eliminate the possibility of off-target effects, an siRNA-resistant myc-tagged version of HELLS was reintroduced to HELLS-depleted cells (Figures 1H, S1G and S1H). U2OS cells transfected with siHELLS and an empty vector displayed persistence of γ-H2AX foci in G2 cells following irradiation relative to cells transfected with control siRNA. In contrast, cells transfected with siHELLS but expressing ectopic siRNA-resistant myc-HELLS did not display elevated numbers of γ-H2AX foci. This confirms that HELLS contributes to efficient repair of IR-induced DSBs in G2.

Disruption of helicase motif I (Walker A box) disrupts ATP binding and remodelling activity in other ATP-dependent chromatin remodelling enzymes (Laurent, Treich and Carlson, 1993; Dong *et al*., 2014). Introduction of an siRNA-resistant version of HELLS containing a substitution in motif I (K254R) failed to restore the number of γ-H2AX foci after 24 hours to the levels observed in cells transfected with either control siRNA or siHELLS complemented with wt HELLS (Figures 1H and S1G). The mutant HELLS protein was expressed at similar levels to the wt protein (Figures S1H); the ability of HELLS to bind ATP and likely remodel chromatin is required for its role in promoting DSB repair.

### HELLS depletion results in defects in the homologous recombination pathway of DSBR

We find that unirradiated HELLS-depleted cells possess elevated numbers of γ-H2AX foci. Spontaneously occurring breaks are predominantly the result of collapsed replication forks that are primarily repaired by the HR pathway either during S-phase or G2. In addition, slow-clearing IR-induced γ-H2AX foci in G2 have been associated with HR (Beucher *et al*., 2009). Furthermore, HELLS expression is highest in CENPF-positive cells (S/G2) when HR is functional, (Figures 2A and S2A, antibody specificity confirmed by knock-down in Figure S2B) and in proliferating tissues active in recombination (Lee *et al*., 2000; Raabe *et al*., 2001). Taken together, this led us to examine whether HELLS participates in the HR pathway of DSBR.

**Figure 2.**
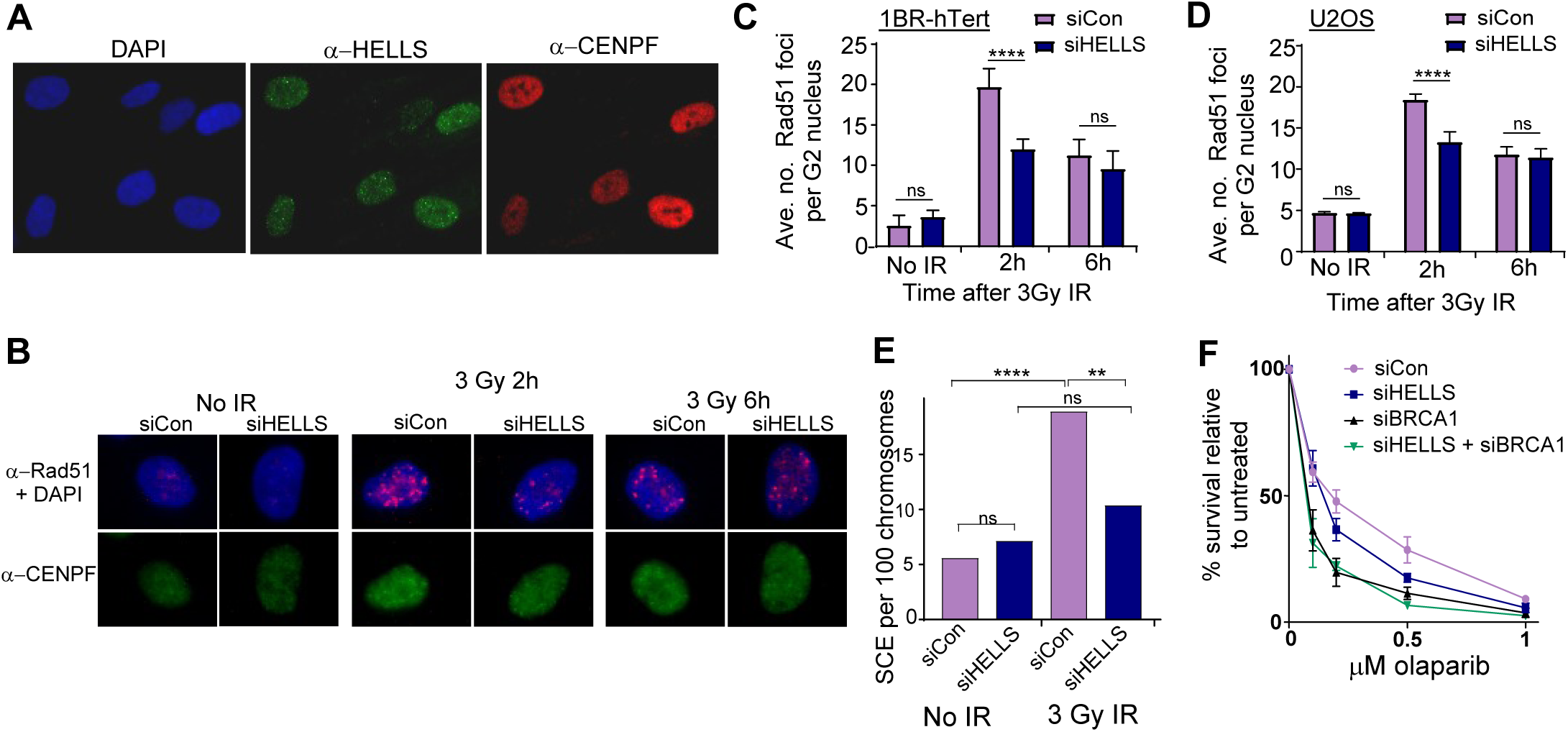
HELLS depletion results in defects in the homologous recombination pathway of DSBR. **(A)**Immunofluorescence images of endogenous HELLS expression and localisation. **(B)** Representative immunofluorescent images of Rad51 foci in G2 cells following IR in 1Br-hTert cells transfected with the indicated siRNA. **(C)** Quantification of IR-induced Rad51 foci in G2 nuclei of 1BR-hTert cells transfected with indicated siRNA. **(D)** Quantification of IR-induced Rad51 foci in G2 nuclei of U2OS cells transfected with indicated siRNA. **(E)** Quantification of sister chromatid exchange events in HeLa cells transfected with the indicated siRNA and analysed 12 h after 3 Gy IR. Frequency of exchange events was calculated per 100 chromosomes from a minimum of 35 spreads from two independent experiments and analysed by an unpaired t-test. **(F)** Viability curves from clonogenic assays of 1Br-hTert cells transfected with the indicated siRNA following exposure to olaparib. **** indicates a p-value of <0.0001, ** indicates p-value of <0.01 and ns indicates not significant by unpaired two-tailed t-test analysis. Data in panels C, D and F are represented as mean +/- sem from three independent experiments.

During G2, the HR-specific RAD51 protein accumulates at IR-induced foci (Figure 2B). Compared with control cells, we observed fewer RAD51 foci 2 hours after IR and a reduction in the percentage of G2 nuclei with more than 20 RAD51 foci in 1BR-hTert cells depleted of HELLS (Figures 2C, S2C and S2D). This defect in RAD51 foci formation was confirmed in a second cell line, U2OS (Figures 2D, S2E, S2F and S2G). In both cell lines, a significant difference was no longer present 6 h after irradiation, when RAD51 foci in control cells have begun to disperse as repair proceeds. These data demonstrate that HELLS facilitates the homologous recombination pathway of DSB repair.

To examine whether the defect in RAD51 accumulation at sites of damage translates to an impact on HR activity, we examined sister chromatid exchange (SCE) events and sensitivity to PARP inhibition. SCEs are the result of HR activity and can be induced by IR or mitomycin c. Metaphase spreads from HeLa cells depleted of HELLS showed fewer IR-induced SCEs than control cells, but no change in the frequency of spontaneous SCEs (Figure 2E). The frequency of SCEs following exposure to mitomycin c was also lower in HELLS depleted cells (Figure S2H and S2I). Again, this is reminiscent of downregulation of ATM or Artemis, which does not affect levels of spontaneous SCEs, but decreases the frequency of SCEs after irradiation (Beucher *et al*., 2009). Another well-documented property of cells with an HR defect is their sensitivity to PARP inhibitors (PARPi) (Bryant *et al*., 2005; Farmer *et al*., 2005; McCabe *et al*., 2006). As established, depletion of the BRCA1 HR protein results in sensitivity to the PARPi, olaparib (Figure 2F). Cells depleted of HELLS also displayed reduced survival on exposure to olaparib, relative to cells transfected with control siRNA (Figure 2F). HELLS-deficient cells were less sensitive to PARPi than BRCA1-deficient cells, suggesting that HELLS does not act as an essential, core HR factor. Co-depletion of BRCA1 and HELLS resulted in a survival frequency indistinguishable from that of BRCA1 depletion alone, indicating an epistatic relationship between HELLS and BRCA1. Taken together, these data suggest that HELLS promotes HR activity.

The murine homologue of HELLS, LSH, has been best-studied for its role in *de novo* methylation of repetitive DNA during development (Yan *et al*., 2003; Myant *et al*., 2011; Yu *et al*., 2014; Samuelsson *et al*., 2016). LSH was found to interact with DNA methyltransferase DNMT3b and indirectly associate with the maintenance DNA methyltransferase, DNMT1 (Myant and Stancheva, 2008). We did not observe a defect in IR-induced RAD51 foci accumulation in cells depleted for DNMT3b or the DNMT1 (Figures S2J-L). Knock-down of DNMT3b and DNMT1 were confirmed by western blot analysis and did not affect HELLS expression (Figures S2M and S2N). Furthermore, knock-down of HELLS did not result in decreased levels of 5-methyl cytosine in MRC5 cells or MEFs within the timescale of our analysis (Suzuki *et al*., 2008; Burrage *et al*., 2012; Ren *et al*., 2019). As such, the function of HELLS in HR appears independent of its role in facilitating DNA methylation.

### HELLS is not a core HR factor but facilitates homologous recombination within heterochromatin

We observed fewer IR-induced Rad51 foci, but not ablation of foci formation in cells depleted of HELLS. This led us to consider whether there may be a subset of breaks that require HELLS for their efficient repair via HR. We utilised the DR-GFP assay to study the effect of HELLS on HR activity (Weinstock *et al*., 2006). In this system, a reporter cassette comprising GFP interrupted by an I-SceI endonuclease site and a downstream fragment of GFP has been integrated into the genome of U2OS cells (Nakanishi *et al*., 2011). Induction of I-SceI results in HR repair of the direct repeats to generate functional GFP (Figure 3A). Depletion of the core HR factor BRCA1, resulted in fewer GFP-positive cells relative to control cells. However, cells depleted of HELLS did not show a significant HR defect using this reporter and downregulation of both HELLS and BRCA1 did not further reduce the number of GFP-positive cells compared to depletion of BRCA1 alone (Fig 3A). This indicates that HELLS is not an essential HR factor. However, if HELLS functions to promote HR within a specific genomic context such as heterochromatin, location of the DR-GFP reporter construct within a euchromatic region of genome would not be expected to result in a defect in this assay. Consistent with this, cells defective in Artemis do not display a defect in the DR-GFP assay and loss of ATM activity results in only a modest decrease in activity (Beucher *et al*., 2009); both ATM and Artemis have been implicated in HR repair of breaks within heterochromatin.

**Figure 3.**
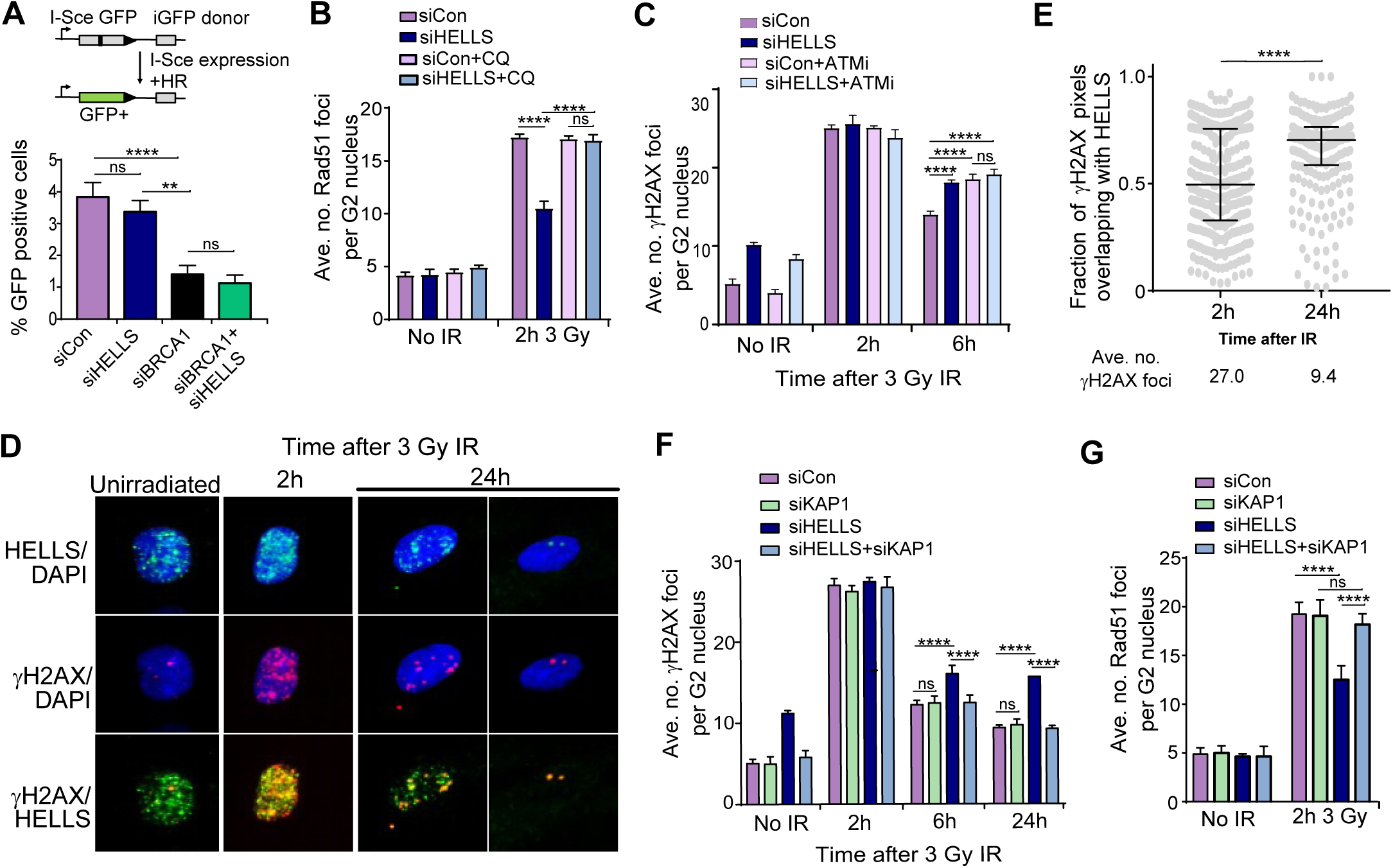
HELLS is not essential for HR but facilitates homologous recombination within heterochromatin. **(A)** HR efficiency assay in U2OS DR-GFP cells transfected with I-SceI expression plasmid and the indicated siRNA. GFP positive cells represent HR repair using a downstream wt-GFP sequence as a donor template and were analysed by FACS. **(B)** Quantification of rescue of Rad51 foci 2 h following 3 Gy irradiation in 1BR-hTert cells transfected with indicated siRNA by the addition of 40 µg/ml chloroquine 2 h prior to irradiation. **(C)** Quantification of clearance of γ-H2AX foci by following 3 Gy irradiation in 1BR-hTert cells treated with indicated siRNA in the absence or presence of 10 µM ATMi, which was added 1 h prior to irradiation. **(D)** Representative immunofluorescent images of HELLS and γ-H2AX foci colocalisation. **(E)** Quantification of colocalisation of γ-H2AX with HELLS at 2 h and 24 h following irradiation with 3 Gy IR. Data represent the fraction of γ-H2AX positive pixels that overlap with HELLS-positive pixels for a minimum of 242 nuclei from three independent experiments, with median value and interquartile range shown. The average number of γ-H2AX remaining from data in Figure 1F is indicated. **(F)** Quantification of rescue of clearance of γ-H2AX foci following 3 Gy irradiation by KAP-1 depletion in 1BR-hTert cells. Data for siCon and siHELLS alone is the same as that shown in Figure 1F and S1E. **(G)** Quantification of rescue of Rad51 foci formation 2 h following 3 Gy irradiation by KAP-1 depletion in 1BR-hTert cells treated with indicated siRNA. **** indicates a p-value of <0.0001, ** indicates p-value of <0.01 and ns indicates not significant by unpaired two-tailed t-test analysis. Data in panels A-C, F and G are shown as the mean +/- sem from three independent experiments.

As HELLS is a member of the Snf2-like family of chromatin remodelling enzymes, we investigated whether relaxation of chromatin structure could overcome the requirement for HELLS in HR repair of IR-induced breaks during G2. We treated cells transfected with control or HELLS siRNA with chloroquine, to yield more open chromatin (Murr *et al*., 2006; Nakamura *et al*., 2011) and analysed IR-induced RAD51 foci formation in G2 (Figures 3B, S3A and S3B). As shown above, downregulation of HELLS resulted in reduced numbers of RAD51 foci 2 hours after irradiation. This defect in RAD51 foci formation was rescued by incubation of HELLS-depleted cells with chloroquine (Figure 3B and S3A), consistent with HELLS being non-essential for HR but facilitating HR in compact chromatin.

Breaks within densely compact heterochromatin require local opening of chromatin structure to be repaired efficiently. During G1, the CHD3 and SMARCA5 chromatin remodellers contribute to repair within heterochromatin (Goodarzi, Kurka and Jeggo, 2011; Costelloe *et al*., 2012; Klement *et al*., 2014). During G2, slow-clearing γ-H2AX foci are proposed to represent breaks within heterochromatic regions of the genome, repaired by HR in an ATM-dependent manner (Goodarzi *et al*., 2008; Beucher *et al*., 2009). Persistent γ-H2AX foci occur in G2 cells depleted of Artemis, RAD51 and BRCA2 or when phosphorylation of the heterochromatin protein KRAB associated protein 1 (KAP-1) is prevented by downregulation of ATM or mutation of the KAP-1 phosphorylation site (Ziv *et al*., 2006; Goodarzi *et al*., 2008). Importantly, depletion of KAP-1 rescues the DSB repair defect of G2 cells downregulated for BRCA2 or ATM (Beucher *et al*., 2009; Geuting, Reul and Löbrich, 2013). Given the association of slow-repairing breaks with HR within heterochromatin, the role of mouse LSH in DNA methylation at heterochromatin (Muegge, 2005; Yu *et al*., 2014; Ren *et al*., 2015) and the requirement for chromatin remodelling enzymes for repair of heterochromatic DSBs during G1, we hypothesised that it may be breaks within heterochromatin that are dependent on HELLS for efficient repair during G2.

Cells treated with ATMi display persistence of IR-induced breaks in G2 which is alleviated by downregulation of the heterochromatin protein KAP-1 (Beucher *et al*., 2009). We find that depletion of HELLS alone is sufficient to cause persistence of γ-H2AX foci, even in the presence of active ATM. We compared the effect of HELLS-depletion to the effect of ATM inhibition on γ-H2AX foci numbers following IR in G2. HELLS-depleted cells and those treated with ATMi possessed similar numbers of unrepaired breaks 6 hours after IR exposure (Fig 3C and S3C). In addition, combining transfection of HELLS siRNA with ATM inhibition did not result in an additive defect, consistent with HELLS and ATM acting epistatically to affect HR repair in heterochromatic regions of the genome in G2.

Immunofluorescent imaging of endogenous HELLS revealed small, bright, nuclear punctae, as well as background pan-nuclear staining in S/G2 cells (Figure 2A). We examined the localisation of γ-H2AX relative to HELLS punctae at different times following irradiation (Figure 3D). HELLS did not form visible IR-induced foci and 2 hours after irradiation γ-H2AX foci were stochastically distributed within the nucleus with no significant colocalisation of HELLS and γ-H2AX i.e. some breaks overlapped with HELLS punctae whilst others did not (Figures 3D and 3E). However, 24 hours after irradiation, when many of the γ-H2AX foci had been resolved due to repair, those γ-H2AX foci that remained showed increased association with HELLS punctae. Analysis of the fraction of γ-H2AX pixels that overlapped with HELLS staining confirmed greater colocalisation of breaks with HELLS 24 hours after irradiation compared to after 2 hours. This suggests that slow-repairing breaks, shown to be associated with heterochromatin, have a greater tendency to overlap with sites enriched in HELLS than faster repairing euchromatic breaks.

Chloroquine relaxes chromatin globally; to specifically examine whether HELLS affects repair within heterochromatin, heterochromatin structure was disrupted by depletion of KAP-1. Downregulation of KAP-1 alone did not alter the number of γ-H2AX foci relative to controls and did not affect the punctate nuclear localisation of HELLS (Figures 3F, S3D and S3E). Strikingly, the elevated numbers of γ-H2AX foci at 6h and 24h post-irradiation in HELLS-depleted cells were alleviated by simultaneous depletion of KAP-1 i.e. persistent breaks in HELLS-deficient cells can be repaired more rapidly if KAP-1 is downregulated. To test whether the HELLS-dependent defect in the HR pathway is also rescued by disruption of heterochromatin, IR-induced RAD51 foci formation was analysed after KAP-1 depletion (Fig 3G and Fig S3F). KAP-1 depletion was able to overcome the reduction in number of RAD51 foci in cells in which HELLS was downregulated (Figure 3G). Within the timeframe of our assays, depletion of HELLS did not result in a significant change in the global levels of the heterochromatin mark H3K9me3 (Figure S3G). These data show that repair can occur in a HELLS-independent manner upon depletion of KAP-1 and support a function for HELLS in relieving or removing a barrier to HR that is posed by heterochromatin.

Phosphorylation of KAP-1 S824 is ATM-dependent and required for initial chromatin relaxation and efficient repair within heterochromatin during G1 (Ziv *et al*., 2006; Goodarzi, Noon and Jeggo, 2009; Noon *et al*., 2010). In G2 cells, pATM is subsequently lost from break sites during resection to favour HR (Geuting, Reul and Löbrich, 2013). We observed that levels of IR-induced S824 phosphorylation of KAP-1 were not altered by depletion of HELLS, either by western blot or when visualised by IF (Figures S3H and S3I). Therefore, global phosphorylation of KAP-1 alone is insufficient to permit efficient repair within heterochromatin; HELLS may either act downstream or perform a distinct function to KAP-1 phosphorylation.

Together our data are consistent with a model where HELLS promotes repair via HR by overcoming a barrier posed by heterochromatic regions of the genome and consequently depletion of HELLS results in repair of these breaks being impeded.

### HELLS facilitates end-resection

Having established that HELLS contributes to repair within heterochromatin, we began to dissect which step of HR is impacted by HELLS downregulation. Chromatin is a known barrier to the first step of HR, end-resection (Adkins *et al*., 2013; Densham *et al*., 2016). The 3’-ssDNA tails that are generated are coated with RPA, which is subsequently replaced by RAD51. As such, the reduction in IR-induced RAD51 foci formation in HELLS-depleted cells could either be the result of a defect in RAD51 loading or further upstream in the HR pathway. Firstly, we studied accumulation of the ssDNA binding protein, RPA, into IR-induced foci in G2 in HELLS-depleted cells (Figure 4A). Analysis of both the average number of RPA foci per G2 nucleus and the percentage of G2 nuclei with more than 20 foci revealed a defect in RPA foci formation in HELLS-depleted cells after irradiation (Figures 4B, 4C and S4A). Decreased RPA foci formation suggests a defect ssDNA formation in HELLS-depleted cells. ssDNA can be monitored more directly using an assay involving BrdU incorporation into DNA prior to irradiation. Following DSB formation, resection results in regions of ssDNA that can be visualised using an anti-BrdU antibody under non-denaturing conditions (Mukherjee, Tomimatsu and Burma, 2015) (Figure S4B). As anticipated, given its role in initiation of end-resection, depletion of CtIP resulted in formation of fewer BrdU foci than in control cells (Figure 4D). Consistent with the defect in RPA foci formation, cells depleted of HELLS also contained fewer ssDNA foci. The decrease in number of BrdU foci following HELLS downregulation is not as pronounced as the defect in CtIP-depleted cells, consistent with HELLS impacting the efficiency of end-resection at a subset of breaks.

**Figure 4.**
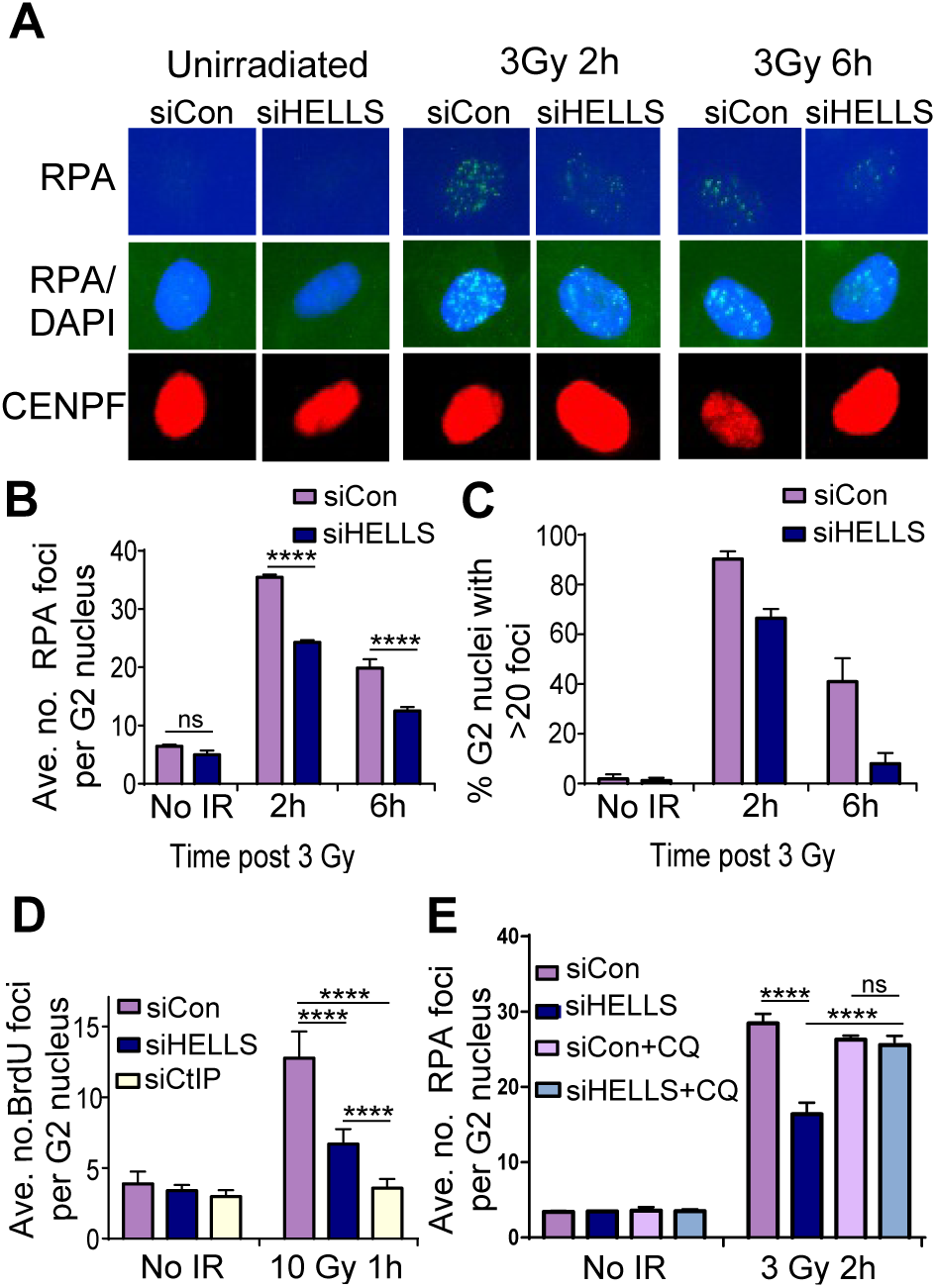
HELLS facilitates end-resection. **(A)** Representative immunofluorescent images of RPA foci in CENPF-positive G2 nuclei following 3 Gy IR. *(B)* Quantification of average number of IR-induced RPA foci per G2 nucleus in 1BR-hTert cells transfected with indicated siRNA. **(C)** Quantification of % of G2 nuclei containing greater than 20 RPA foci in 1BR-hTert cells transfected with siCon or siHELLS. **(D)** Quantification of number of BrdU foci per nucleus 1 h after 10 Gy, visualised under non-denaturing conditions and therefore representing regions of ssDNA, in U2OS cells transfected with indicated siRNA. **(E)** Quantification of rescue of RPA foci 2 h following 3 Gy irradiation in 1BR-hTert cells treated with indicated siRNA by the addition of 40 µg/ml chloroquine 2 h prior to irradiation. **** indicates a p-value of <0.0001, and ns indicates not significant by unpaired two-tailed t-test analysis. Data in panels B-E are shown as the mean +/- sem from three independent experiments.

Using IR-induced RPA foci number as a marker of ssDNA formation, we examined whether relaxation of chromatin structure could also rescue the end-resection defect of HELLS-depleted cells. Addition of chloroquine restored the number of IR-induced RPA foci in HELLS-depleted cells to the levels observed in cells transfected with control siRNA (Figure 4E). We find that HELLS facilitates end-resection, the initial step of the HR pathway, within heterochromatin.

During G2, 53BP1 both negatively and positively influences HR. In G2 cells a BRCA1-dependent mechanism alleviates the inhibitory effect of 53BP1 on resection by promoting 53BP1 dephosphorylation but 53BP1 also contributes to HR via relaxation of densely packed heterochromatin (Kakarougkas *et al*., 2013). We examined epistasis of HELLS and 53BP1 in ssDNA BrdU foci and RPA foci accumulation. In both assays, 53BP1 depletion failed to rescue the end-resection defects observed in HELLS depleted cells (Figures S4C and S4D). This indicates that the mechanism by which HELLS facilitates end-resection is not through removing the inhibitory effect of 53BP1. Indeed, a small additive defect was observed when HELLS and 53BP1 were simultaneously downregulated, indicating that they do not exclusively function in the same pathway.

There is precedent for involvement of ATP-dependent chromatin remodellers promoting end-resection during DSB repair. Phosphorylated SMARCAD1 is recruited to breaks where it acts to facilitate long-range end-resection by repositioning 53BP1 (unlike HELLS-depleted cells, the resection defect in SMARCAD1-depleted cells is rescued by downregulation of 53BP1) (Costelloe *et al*., 2012; Densham *et al*., 2016). We examined epistasis between SMARCAD1 and HELLS in ssDNA accumulation and found that SMARCAD1 depleted cells displayed a more severe defect than those depleted of HELLS (Fig S4E). Co-depletion of SMARCAD1 and HELLS reduced ssDNA formation to an extent comparable, or slightly greater than that observed upon SMARCAD1 downregulation. We conclude that HELLS and SMARCAD1 serve non-redundant functions in facilitating resection. This highlights the different chromatin remodelling requirements at different steps in DSB repair and in different chromatin contexts.

### The HELLS protein interacts with CtIP and contributes to its accumulation at IR-induced breaks

Chromatin structure may affect recruitment and retention of the end-resection machinery and/or influence the processivity or efficiency of nuclease activity. To examine if accumulation of proteins involved in the initial steps of end-resection was HELLS-dependent, we introduced a GFP-tagged version of CtIP to U2OS cells and analysed its accumulation at IRIF. Four hours after exposure to IR, cells depleted of HELLS displayed fewer nuclear GFP-CtIP foci than control cells (Figures 5A, 5B and S5A). Therefore, HELLS appears to enable either recruitment or retention of the CtIP protein at breaks, in accordance with a function in promoting initiation of end-resection. There was not a reciprocal effect on localisation, as the punctate nuclear localisation of HELLS was not grossly altered by depletion of CtIP (Figure 5C); while HELLS impacts accumulation of CtIP, localisation of HELLS is independent of CtIP.

**Figure 5.**
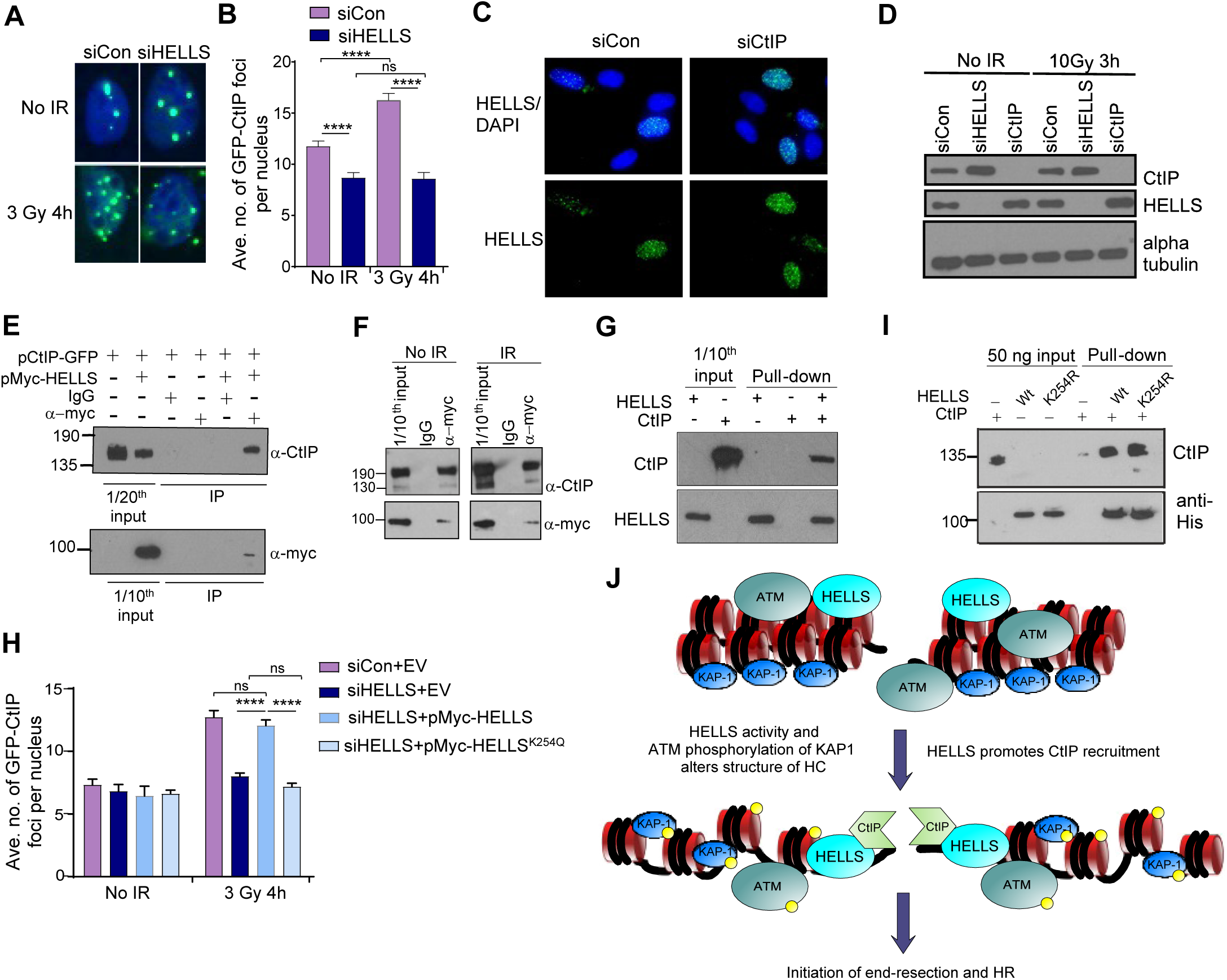
HELLS interacts with CtIP and contributes to its accumulation at IR-induced breaks. **(A)** Representative immunofluorescent images of CtIP-GFP foci 4h following 3 Gy IR in U2OS cells transfected with CtIP-GFP expressing plasmid and the indicated siRNA. **(B)** Quantification of average number of IR-induced CtIP-GFP foci in U2OS cells transfected with CtIP-GFP expressing plasmid and indicated siRNA. **(C)** Representative immunofluorescent images showing that the punctate localisation of HELLS in 1BR-hTert nuclei is not disrupted by depletion of CtIP. **(D)** Western blot analysis of levels of CtIP and HELLS in whole cell extracts prepared from U2OS cells transfected with the indicated siRNA. Alpha tubulin was used as a loading control. **(E)** Immunoprecipitation of CtIP with Myc-HELLS from nuclear extracts of U2OS cells transfected with indicated plasmids, analysed by western blotting. **(F)** Immunoprecipitation of CtIP with Myc-HELLS from nuclear extracts of U2OS cells transfected with indicated plasmids 1 h following 3 Gy IR, analysed by western blotting. **(G)** Western blot analysis of pull-down assay using nickel beads and purified recombinant His-HELLS and CtIP proteins. **(H)** Quantification of rescue of average number of IR-induced CtIP-GFP foci in U2OS cells transfected with CtIP-GFP expressing plasmid, HELLS siRNA and constructs expressing siRNA-resistant wt or ATPase mutant HELLS. For panels B and I, foci were counted blind and data represent the mean +/- sem from three independent experiments. **** indicates a p-value of <0.0001, and ns indicates not significant by unpaired two-tailed t-test analysis. **(I)** Western blot analysis of pull-down of CtIP by wt and K254R mutant HELLS proteins using anti-His and protein G Dynabeads. **(J)** Cartoon depicting model of the role of HELLS at a double-strand break within heterochromatin.HELLS activity along with ATM-dependent phosphorylation of KAP-1 results in changes to the structure of heterochromatin that facilitate accumulation of CtIP. CtIP recruitment or retention is promoted by interaction with HELLS in order to help initiate end-resection and HR.

Levels of CtIP protein, evaluated by western blotting of whole cell extracts, are not decreased (in fact they are modestly elevated) upon depletion of HELLS, indicating that the defect in formation of IR-induced CtIP foci is not the result of reduced CtIP expression (Figure 5D). Nor is there a significant effect of CtIP depletion on HELLS levels or significant changes in HELLS expression following irradiation.

One mechanism by which HELLS could promote accumulation of CtIP at IR-induced foci is through interaction of HELLS and CtIP proteins. We introduced to U2OS cells a plasmid expressing CtIP-GFP either by itself or along with a plasmid expressing Myc-HELLS. Immunoprecipitation using an anti-myc antibody from cells expressing myc-tagged HELLS, resulted in the coimmunoprecipitation of CtIP-GFP (Figure 5E). Immunoprecipitation of CtIP-GFP with Myc-HELLS was not stimulated by irradiation (Figure 5F). An interaction between HELLS and CtIP proteins was confirmed using recombinant purified proteins. CtIP and His-tagged human HELLS were expressed and purified from insect cells (Figure S5B) and His-HELLS immobilised on cobalt beads pulled-down CtIP (Figure 5G). We did not observe any stimulation of the ATPase activity of HELLS by CtIP *in vitro* (data not shown). These data demonstrate a direct protein-protein interaction between HELLS and CtIP, which may contribute to the HELLS-dependent accumulation of CtIP at IR-induced breaks during G2 and subsequent end-resection.

CtIP foci accumulation after IR was rescued by expression of an siRNA resistant version of HELLS, wt Myc-HELLS, in cells in which endogenous HELLS had been depleted. This confirms that HELLS function is required for CtIP localisation following IR (Figure 5H and S5C). Introduction of an siRNA-resistant ATP-binding site mutant of HELLS did not result in rescue of CtIP IRIF foci formation (HELLS K254R retains the ability to associates with the chromatin fraction Figure S5D). Recombinant HELLS K254R protein was still able to bind to CtIP (Figure 5I), suggesting that the ability of HELLS to remodel chromatin structure, as well as bind CtIP contributes to full stimulation of initiation of end-resection.

## DISCUSSION

This work uncovers a role for chromatin remodelling activity in DSBR repair in G2 and provides details of the function of HELLS in maintenance of genome stability. We find that the human HELLS chromatin remodelling enzyme, is required for efficient repair of two-ended DSBs within heterochromatin in G2. Repair of these heterochromatic breaks by HR is impaired and end-resection defective when HELLS is downregulated, with the ATP binding site of HELLS required for its function in DSBR. We demonstrate that HELLS facilitates accumulation of CtIP at IR-induced breaks and identify an interaction between HELLS and CtIP proteins. Compact heterochromatin poses a particular challenge to DNA repair processes and our findings highlight the importance of specific chromatin remodelling activities during repair in different regions of the genome in order to prevent genome instability.

The structure of heterochromatin acts as a barrier to recruitment and activity of DSB repair proteins. We propose that HELLS establishes a chromatin structure that permits recruitment, retention and activity of the end-resection machinery at breaks within heterochromatin. Given the requirement for an intact ATPase motif within HELLS, and that disruption of heterochromatin structure by depletion of KAP-1 rescues repair defects that occur upon HELLS downregulation, this likely involves decondensation or relaxation of heterochromatin, stimulated by chromatin remodelling by HELLS. We propose that HELLS facilitates end-resection through a combination of chromatin remodelling to generate a chromatin environment suitable for recruitment of the end-resection machinery, and interaction of HELLS and CtIP to assist recruitment or retention of resection proteins (Figure 5J). Given that depletion of KAP-1 rescues the resection defect of HELLS-depleted cells, and that persistent breaks display a tendency to colocalise with HELLS, we favour a model where the contribution of HELLS to DSB repair is direct, rather than due to misregulation of transcription. Consistent with this, transcription of known DNA repair genes is not significantly changed and only 0.4 % of transcripts are altered in LSH -/- MEFs (Burrage *et al*., 2012).

We did not detect a change in the amount of chromatin-bound HELLS after IR and the pattern of HELLS localisation by immunofluorescence was indistinguishable from that of undamaged cells. Therefore, HELLS may be constitutively present at heterochromatin and its remodelling activity stimulated by break formation or alternatively, post-translational modification of either HELLS or CtIP could regulate interaction with the end-resection machinery. The question of how the HELLS-CtIP interaction is regulated will be of future interest.

We find that HELLS is not an essential global HR factor; HELLS-deficient cells did not show a significant loss of HR activity in a frequently utilised direct repeat reporter assay, disruption of chromatin structure can rescue RAD51 and RPA foci formation in cells downregulated in HELLS, and olaparib sensitivity is more pronounced following depletion of the core HR protein BRCA1 is downregulated than HELLS. However, consistent with a function in enabling HR, HELLS expression is highest in proliferating tissues, active in recombination such as testes and thymus. We uncovered impaired accumulation of HR pathway proteins in HELLS downregulated cells, but also defects in some readouts of HR repair. In agreement with the PARPi sensitivity that we observed, the HELLS gene was identified in a genome-wide shRNA screen for olaparib sensitivity, and as part of a gene signature associated with response to PARP inhibition in a panel of triple-negative breast cancer cell lines (Bajrami *et al*., 2014; Hassan *et al*., 2017). Rather that acting as a core HR protein, we suggest that HELLS facilitates recombination at a subset of DSBs, namely those located within heterochromatin.

In support of a function at heterochromatin, LSH colocalises with chromocenters and HP1 (Yan *et al*., 2003) and was also found to ChIP (although not exclusively) to repetitive heterochromatin-enriched regions of the genome (Von Eyss *et al*., 2012), and colocalise with late replication foci (Yan *et al*., 2003). Notably, HELLS mutations have been identified in ICF syndrome, which is characterised by genomic rearrangements in heterochromatic pericentromeric regions (Thijssen *et al*., 2015). The loss of functional HELLS and efficient HR repair of breaks within heterochromatin may contribute to the pericentric instability observed in cells from these patients

Phosphorylation of the KAP-1 heterochromatin protein is necessary for efficient repair of heterochromatin breaks in G2. As both HELLS and KAP-1 phosphorylation are epistatic with ATM, HELLS could act downstream of KAP-1 phosphorylation (KAP-1 phosphorylation still occurs on HELLS downregulation). Alternatively, the effect of HELLS on heterochromatin structure during DSBR may be distinct from remodelling events that occur upon KAP-1 phosphorylation. We cannot exclude that HELLS may affect a second, downstream step of repair within heterochromatin, in addition to recruitment of the resection machinery. For example, HELLS could contribute to efficient progression of resection, recruitment of HC factors that occurs following initial local chromatin decompaction or to relocalisation of breaks within the nucleus (Geuting, Reul and Löbrich, 2013; Alagoz *et al*., 2015; Tsouroula *et al*., 2016).

In contrast to cells depleted of HELLS or BRCA2 or treated with ATMi, cells depleted of CtIP do not show persistent G2 γ-H2AX foci following IR (Shibata *et al*., 2011; Geuting, Reul and Löbrich, 2013). In CtIP-depleted cells, resection is not initiated and therefore breaks within heterochromatin that would normally be repaired by HR can be redirected into an end-joining pathway. Despite displaying a defect in CtIP accumulation in response to IR, HELLS-depleted cells possessed elevated numbers of unrepaired breaks at later time points after irradiation, suggesting that an alternative repair pathway is also unable to function efficiently; co-depletion of HELLS and CtIP resulted in the same persistence of γ-H2AX foci as depletion of HELLS alone (Figures S5E and S5F). This indicates that the chromatin remodelling activity of HELLS is required not only for the preferred HR pathway, but also for factors in the compensating end-joining pathway to access heterochromatin during G2. In this study we have examined HR repair of IR-induced breaks in G2, however breaks within heterochromatin are also repaired with slow kinetics in G1 (although in this instance they are repaired by NHEJ). Recently, resection-dependent c-NHEJ involving CtIP has been shown to occur in G1 (Shibata *et al*., 2017) and it will be of interest to examine if HELLS additionally impacts repair by NHEJ within heterochromatin during G0/G1 or compensatory end-joining when resection is inhibited during G2.

We observed elevated spontaneous H2AX phosphorylation and persistence of IR-induced γ-H2AX foci in HELLS-deficient 1BR-hTert immortalised human fibroblasts. It has previously been reported that mammalian HELLS promotes efficient phosphorylation of H2AX (Burrage *et al*., 2012). There are several possible explanations for this apparent discrepancy including cell cycle effects, our use of lower, non-lethal doses of radiation, x-rays vs γ-gays and method of HELLS-depletion. There may also be differences between murine and human DNA damage responses within heterochromatin. Consistent with our data, spontaneous γ-H2AX foci have recently been observed in an undamaged HEK293 HELLS KO cell line (Unoki *et al*., 2018).

HELLS/LSH have been most extensively studied for their role in *de novo* DNA methylation. We did not observe HR pathway defects upon depletion of DNMT3b or DNMT1 DNA methyltransferases and in agreement with this, ICF patient cells carrying mutant DNMT3b display unaltered DSBR repair (Brunton *et al*., 2011). Nonetheless, DNA methylation may still contribute to the overall decreased genome stability in cells lacking functional HELLS; both defective DSBR and hypomethylation and misfunction of centromeres during chromosome segregation could result in increased incidence of micronuclei. This may be relevant to ICF syndrome patients with HELLS mutations, given that hypomethylation of repeat sequences is observed at a subset of chromosomes and the most frequently observed mutations in ICF syndrome are within DNMT3b (Hansen *et al*., 1999; Jin *et al*., 2008). HELLS has been associated with hypomethylation of specific repeat elements in some colorectal tumours (Samuelsson *et al*., 2016). However, DNA methylation does not occur in budding yeast and consequently it seems likely to be the function at heterochromatic regions or DNA repair that is conserved in the yeast homolog, Irc5.

Many chromatin remodelling enzymes function in several cellular processes, including DNA repair, transcription, chromosome segregation and replication. These pleiotropic effects are expected to contribute to the prevalence of mutations or misregulation of chromatin remodelling enzyme subunits in cancer. HELLS may perform multiple roles within the cell, such that HELLS downregulation or mutation may increase genetic instability and additional effects on DNA methylation and transcription could contribute to tumourigenesis. By elucidating a mechanism through which HELLS impacts DSB repair, this study raises the potential for targeted therapeutic treatment such as PARP inhibitors, if tumours could be stratified by their HELLS status. Notably, HELLS overexpression is associated with progression of squamous cell, oropharyngeal and prostate cancers, with high HELLS levels also found in bladder, retinal, lung and ovarian tumours and HELLS expression a potential biomarker for melanoma metastasis (Kim *et al*., 2010; Janus *et al*., 2011; Keyes *et al*., 2011; Von Eyss *et al*., 2012). It will be of interest to determine whether HELLS upregulation also affects genome stability or if its impact on cancer progression is due to another mechanism such as transcriptional misregulation of genes that drive proliferation. In either case, it appears that there is an optimal level of HELLS activity for preventing tumourigenesis.

## MATERIALS AND METHODS

### Cell culture and irradiation

1BR-hTERT human fibroblasts (provided by Prof P Jeggo), U2OS and HeLa cells were cultured in Dulbecco’s modified Eagle’s medium (D-MEM) GlutaMAX™ (Gibco) supplemented with 10% foetal bovine serum (FBS)(Gibco), 100 units/ml penicillin, and 100 μg/ml streptomycin (Gibco). Cells were grown at 37°C, 95% humidity, 5% CO_2_. Cells were irradiated 24 h after siRNA knock-down, with a Cs137 source RX30/55M Irradiator (Gravatom Industries Ltd).

For G2 analyses, 3 µg/ml aphidicolin (Sigma-Aldrich) was added prior to IR, to prevent progression of S-phase cells into G2. Where indicated, 10 µM ATM inhibitor (Ku-55933) was added 1 h prior to IR or 40 µg/ml chloroquine (Sigma-Aldrich) 2 h prior to irradiation.

### siRNA knock-downs, plasmids and transfection

siRNA oligonucleotides were transfected using Lipofectamine RNAiMAX reagent (Invitrogen). 5 nM siRNA duplexes were used for reverse transfection of 5 × 10^5^ of logarithmically growing cells in a 6 well plate. siRNA sequences are provided in the appendix. For complementation experiments, knock-down was performed using a single HELLS siRNA oligonucleotide 24h prior to transfection of pMyc-HELLS plasmid DNA using Lipofectamine 3000 (Invitrogen). Media was changed 6 h after plasmid transfection and cells incubated for 24h prior to irradiation. pMyc-HELLS expresses N-terminally Myc-tagged siRNA-resistant HELLS from a CMV promoter. It was constructed by introducing a myc-epitope and MCS downstream of the CMV promoter of pUHD15-1 (lacking the tTA), to create pALC43 (sequence available on request). HELLS cDNA (from NM_18063.3 in pReceiver-M98 from Tebu) into which mutations that confer resistance to siHELLS were introduced (ATAGGGAGAGCACAG) was then cloned into the NotI-SalI sites of pALC43 to create pMyc-HELLS.

### Cell extracts

For whole cell extracts, cells were lysed into RIPA buffer (50 mM Tris pH 7.5, 150 mM NaCl, 5 mM EDTA, 10 mM K_2_HPO_4_, 10% (v/v) glycerol, 1% (v/v) Triton X-100, 0.05% SDS, 1mM DTT, protease inhibitors (Roche) and debris removed prior to loading. For nuclear extracts, cells were resuspended in hypotonic buffer (100 mM HEPES pH 7.9, 10 mM KCl, 1 mM MgCl_2_, 1mM DTT, 1% TritonX-100 and protease inhibitors (Roche)) and sheared following addition of 0.5% NP-40. Nuclei were pelleted by centrifugation at 3000 rpm for 5 min at 4°C and were resuspended in hypotonic buffer containing 1U/µl of benzonase (Merck). Extracts were incubated on ice for 30 min, before addition of 400mM NaCl and incubation for a further 1h at 4°C, prior to centrifugation at 13000 rpm for 15 min at 4°C and using the supernatant for analysis. For chromatin extracts, nuclei were harvested as above and the nuclear pellet resuspended in 100 µl extraction buffer (3mM EDTA, 0.2mM EGTA and protease inhibitors (Roche)). Samples were spun at 13000 rpm for 5 min at 4°C and the chromatin pellet resuspended in 20 µl 0.2 M HCl before neutralisation with 80 µl 1 M Tris-HCl pH 8.

### Immunofluorescence and colocalisation analysis

Cells grown on glass coverslips were pre-extracted with 0.2% Triton-X 100 in PBS for 1 min, before fixation with 4% (w/v) paraformaldehyde for 10 min at room temperature. Cells were washed with PBS and incubated with primary antibody diluted in 2% (w/v) bovine serum albumin (BSA) in PBS for 1h at RT. After washing with PBS, cells were incubated with secondary antibody diluted in 2% BSA for 1h at RT, followed by 10 min at RT in 100ng/µl DAPI. After washing in PBS, coverslips were mounted with Vectashield (Vector Laboratories). Antibodies used are provided in the appendix.

For BrdU foci analysis of resection, cells were incubated with 10µg/ml BrdU for 16h and irradiated with 10Gy IR. 1h later, cells were washed with PBS and incubated in 1ml extraction buffer 1 (10mM PIPES pH 7, 100mM NaCl, 300mM Sucrose, 3mM MgCl_2_, 1mM EGTA, 0.5% Triton-X 100) for 10 min on ice. After washing with PBS, cells were incubated with 1ml extraction buffer 2 (10mM Tris-HCl pH 7.5, 10mM NaCl, 3mM MgCl_2_, 1% Tween20, 0.5% sodium deoxycholate) for 10min on ice. After another wash, cells were fixed with paraformaldehyde (4%, w/v) for 20 min on ice, washed in PBS and permeabilised with 0.5% Triton-X 100 in PBS for 10 min on ice. After washing, cells were incubated with 0.5 ml 5% BSA (w/v) in PBS for 20 min on ice and then were incubated overnight with primary antibody for 16h at 4°C and secondary antibody as described above.

For colocalisation studies, cells were fixed with ice-cold methanol for 20 min. Dried slides were washed with PBS and primary and secondary antibody staining was performed as described above. CellProfiler was used to quantify colocalization by measuring the fraction of γ-H2AX positive pixels that overlap with HELLS-positive pixels.

All images were taken using either Leica DMI600 microscope, with Leica DFC365FX monochrome CCD camera, 40x lenses (serial number 506201), acquisition software Leica LAS-X, or using Leica DMR, with R6 Retiga camera (QImaging) and CoolLED pE-300 light source, 40x lenses (serial number 506144), acquisition software: Micro-Manager.

### HR reporter assay

siRNA oligonucleotides were transfected into U2OS-DR-GFP cells (kindly provided by Maria Jasin). 24h later cells were transfected with pCBAS plasmid (expressing I-SceI) and after a further 48 h, cells were harvested. Cells were washed with PBS and analysed for GFP fluorescence by FACS as described in the appendix.

### Clonogenic survival assay

48 h after siRNA transfection, 500 cells were seeded into 6 cm dishes and olaparib (Ku-0059436, Stratech) added at indicated concentrations. Cells were fixed with ice-cold methanol after 14 days and stained with 1% (w/v) crystal violet.

### Sister chromatid exchange assays

HeLa cells transfected with siRNA oligonucleotides were incubated with 10 µM BrdU for 48h. To enrich for mitotic cells, 1 mM caffeine and 0.2µg/ml colcemid were added 8 hours after irradiation with 3 Gy and cells were harvested after a further 4 h. For chromosome spreads, cells were resuspended in 75mM KCl for 16 min at 37°C, before centrifugation at 1500 rpm for 10 min at 4°C. Cells were fixed in 3:1 methanol: glacial acetic acid and metaphases dropped onto slides. Chromosome spreads were airdried, incubated with 10 µg/ml Hoechst, washed with Sorensen buffer pH 6.8, covered with Sorensen buffer and a cover slip and irradiated under 365nm UV lamp for 1h. Coverslips were removed and slides incubated in SSC buffer (2M NaCl, 0.3 M Na-citrate pH 7) at 67°C for 1h before staining with 10% Giemsa (Acros) in Sorensen buffer for 30 min at RT. Slides were washed with water and dried overnight before mounting.

### Co-immunoprecipitation

U2OS cells transfected with the indicated plasmids were irradiated, where shown, with 3 Gy IR 1 hour prior to being harvested. Cells were lysed and nuclear extracts prepared as above, with the exception that NaCl was adjusted to 200 mM. Extracts were incubated for 2 hours with 4 µg α-myc or IgG before addition of Protein G Dynabeads and a further 1 hour incubation at 4°C. 3 × 1ml washes in lysis buffer + 200 mM NaCl were performed and bound protein eluted by boiling prior to analysis by western blotting.

### *In vitro* pull-down assay

Details of purification of recombinant His-HELLS and CtIP proteins are provided in the appendix.

His-select Cobalt affinity beads (Sigma) were equilibrated in binding buffer (40 mM HEPES pH 7.5, 30 mM imidazole pH 8.0, 150 mM NaCl, 0.1% Tween, 10% glycerol, 0.2% BSA) and 100 ng His-HELLS protein bound for 1 h at 4°C. 100 ng recombinant CtIP protein was added and reactions were incubated for 1 hour at 4°C. Alternatively 2 µg His-HELLS and 0.6 µg CtIP were mixed and incubated for 30 min at 4°C before capture using 1 µl anti-His antibody and protein G Dynabeads for 40 min at 4°C. Beads were washed in equilibration buffer and bound protein eluted by boiling in SDS-PAGE sample buffer for analysis by western blotting.

## ACKNOWLEDGEMENTS

We thank Oliver Wilkinson and Mark Dillingham for recombinant CtIP protein, Maria Jasin for U2OS DR-GFP cell line, Stephen Cross and the Elizabeth Blackwell Institute, and its Wellcome Trust ISSF Award for assistance with colocalisation analysis, Sophie Wells for assistance with plasmid construction, members of the DNA-protein interactions unit for helpful discussions and the Bristol Wolfson Bioimaging and FACS Facilities. GK, CET and ALC are supported by Cancer Research UK C49963/A1750.

## AUTHOR CONTRIBUTIONS

Experiments were conceived and designed by GK and ALC and performed by GK, CET, EPS and ALC. GK, ALC analysed data and wrote the manuscript.

---

**Figure S1.**
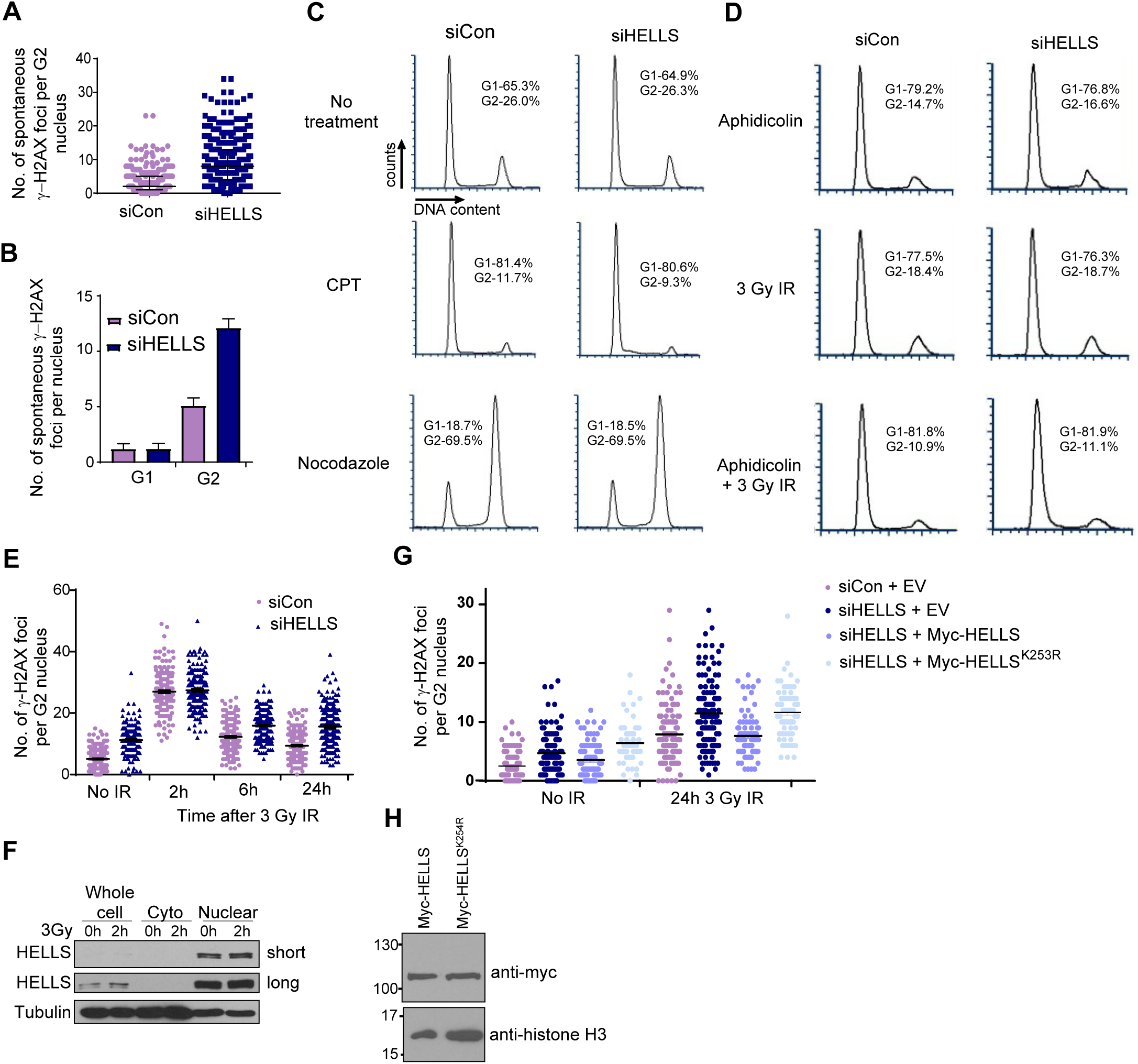
HELLS promotes genome stability in undamaged cells and following IR. **(A)** Counts of spontaneous γ-H2AX foci per G2 nucleus in 1BR-hTert cells transfected with siCon or siHELLS. Data from six independent experiments with median and interquartile range shown. **(B)** Ave. number of spontaneous γ-H2AX foci in G0/G1 serum-starved (5 independent experiments) or G2 (data as Fig1C) nuclei in 1BR-hTert cells transfected with siCon or siHELLS. **(C)** Cell cycle and checkpoint analysis by FACS of 1BR-hTert cells transfected with indicated siRNA. 0.5 µM CPT or 100 ng/ml nocodozole were added at time of siRNA transfection and samples collected after 24 hours. **(D)** Cell cycle analysis by FACS of 1BR-hTert cells transfected with the indicated siRNA 24 h prior to irradiation. 6 µM aphidicolin was added at the time of irradiation and samples collected 24 h after irradiation. **(E)** γ-H2AX foci counts per G2 nucleus in 1BR-hTert cells transfected with siCon or siHELLS at indicated time points following 3 Gy IR. Data is combined from three independent experiments and mean +/- sem shown. **(F)** Western blot analysis of distribution of HELLS in cytoplasmic and nuclear fractions prepared from undamaged 1BR-hTert cells, or 2 h following 3 Gy IR. Alpha tubulin is used as a loading control. **(G)** Counts of the number of IR-induced γ-H2AX foci per nucleus in U2OS cells transfected with siCon or HELLS siRNA and constructs expressing siRNA-resistant wt or ATPase mutant HELLS. Combined data from at least three independent experiments and mean +/- sem shown. **(H)** Western blot analysis of myc-HELLS wt and Myc-HELLS^K254R^ mutant proteins in extracts prepared from U2OS cells transfected with indicated expression plasmid.

**Figure S2.**
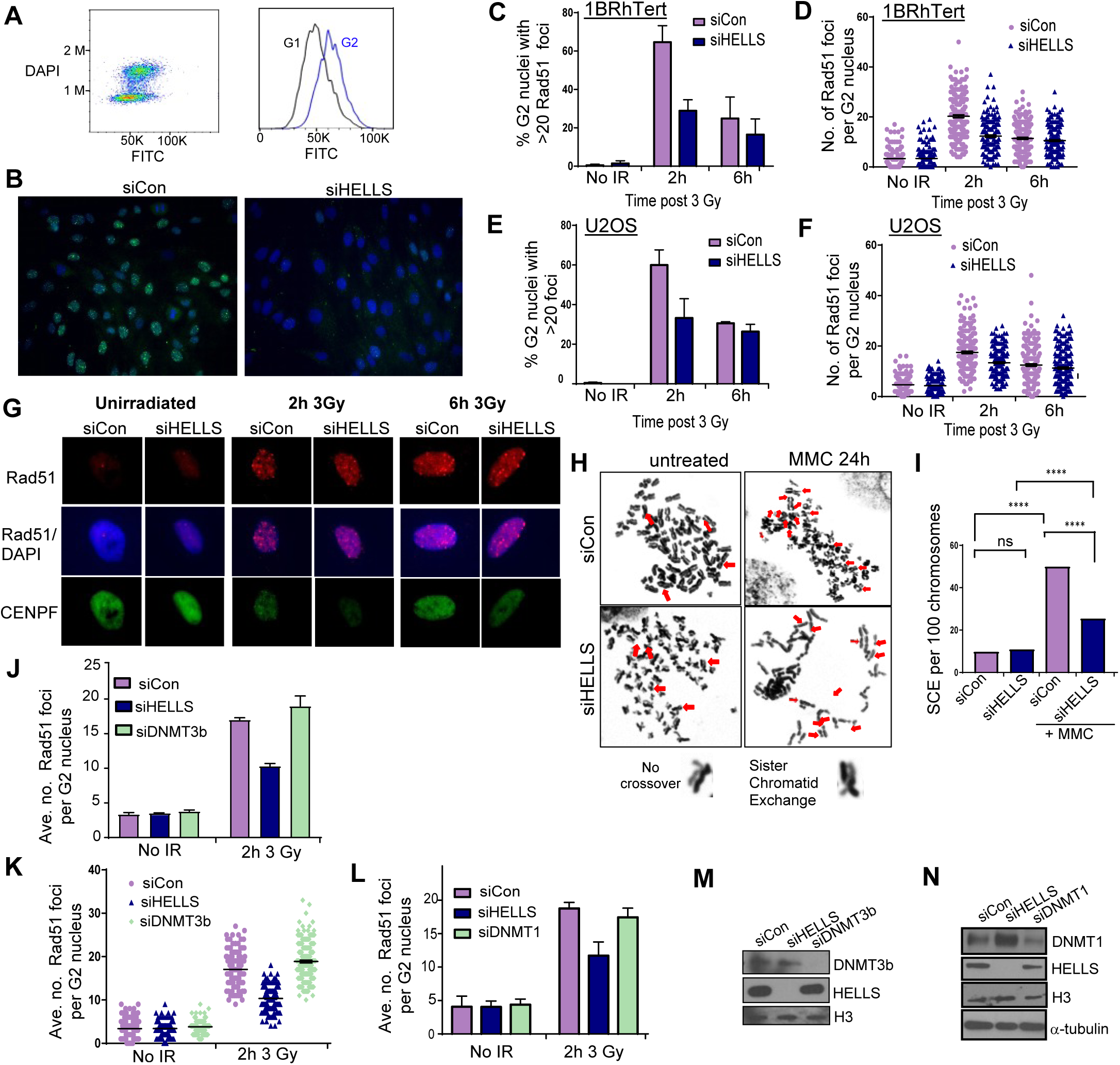
HELLS depletion results in defects in the homologous recombination pathway of DSBR. **(A)** FACS analysis of HELLS expression in G1 vs G2 using anti-HELLS (FITC) and DAPI staining. **(B)** Anti-HELLS and DAPI immunofluorescent images of 1BR-hTert cells transfected with siCon or siHELLS. **(C)** % of G2 nuclei with greater than 20 Rad51 foci in 1BR-hTert cells transfected with siCon or siHELLS. (**D)** Number of Rad51 foci per G2 nucleus in 1BR-hTert cells transfected with siCon or siHELLS **(E)** Quantification of IR-induced Rad51 foci in G2 nuclei of U2OS cells transfected with indicated siRNA. **(F)** Number of Rad51 foci per G2 nucleus in U2OS cells transfected with siCon or siHELLS **(G)** Representative immunofluorescent images of Rad51 foci in U2OS cells transfected with the indicated siRNA following 3Gy IR. **(H)** Representative images of chromosome spreads to analyse SCEs in HeLa cells transfected with siCon or siHELLS following exposure to mitomycin c. **(I)** Frequency of sister chromatid exchange events per 100 chromosomes in HeLa cells transfected with the indicated siRNA and exposed to mitomycin c. Analysis of ∼ 20 metaphases, statistical analyses by unpaired t-test, **** represents p-value <0.0001. **(J)** Quantification of IR-induced Rad51 foci in G2 nuclei of 1BR-hTert cells transfected with indicated siRNA. **(K)** Number of Rad51 foci per G2 nucleus in 1BR-hTert cells transfected indicated siRNA. **(L)** Quantification of IR-induced Rad51 foci in G2 nuclei of 1Br-hTert cells transfected with indicated siRNA. **(M)** Western blot analysis of efficiency of DNMT3b and HELLS depletion by siRNA from nuclear extracts prepared from 1BR-hTert cells. Histone H3 used as a loading control. **(N)** Western blot analysis of efficiency of DNMT1 and HELLS depletion by siRNA from nuclear extracts prepared from 1BR-hTert cells. Alpha tubulin used as a loading control. For panels C,E, J and L, data shown are mean +/sem from three independent experiments.

**Figure S3.**
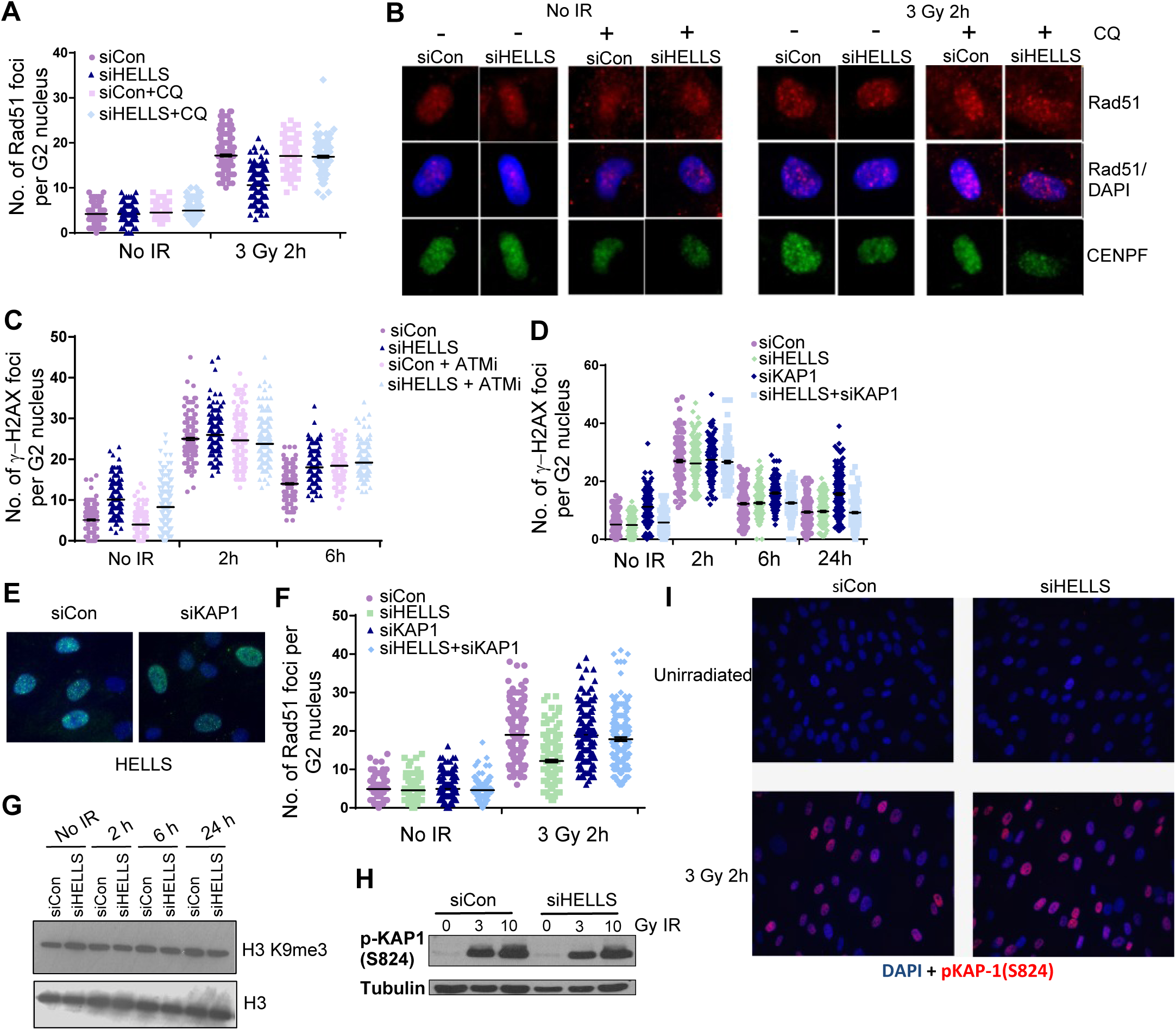
HELLS facilitates homologous recombination within heterochromatin. **(A)**Individual counts of number of spontaneous Rad51 foci per G2 nucleus in 1BR-hTert cells transfected with siCon or siHELLS in the presence or absence of chloroquine 2 h after 3 Gy IR. **(B)** Representative immunofluorescent images of Rad51 foci in 1BR-hTert cells transfected the indicated siRNA with or without addition of 40 µg/ml chloroquine 2 h prior to irradiation. **(C)** Individual counts of number of γ-H2AX foci per G2 nucleus in 1BR-hTert cells transfected with siCon or siHELLS. Where indicated, 10 µM ATMi was added 1 h prior to irradiation. **(D)** Individual counts of rescue of clearance of γ-H2AX foci following 3 Gy irradiation by KAP-1 depletion in 1BR-hTert cells. Data for siCon and siHELLS is the same as that shown in Figure 1F and S1E. **(E)** Representative immunofluorescent images of 1BR-hTert cells indicating that the punctate nuclear localisation of HELLS is not affected by KAP-1 depletion. **(F)** Individual counts Rad51 foci formation per nucleus following 3 Gy irradiation in 1BR-hTert cells transfected with the indicated siRNA. **(G)** Western blot analysis of H3 K9me3 in 1BR-hTert cells transfected with siCon or siHELLS, at indicated time points following 3 Gy IR. Total H3 is used as a loading control. **(H)** Western blot analysis of phosphorylated (S824) KAP-1 in 1BR-hTert cells transfected with siCon or siHELLS 30 min following 3Gy or 10Gy IR. **(I)** Immunofluorescent images of pKAP-1 (S824) in 1BR-hTert cells transfected with siCon or siHELLS, before or 2 h following 3 Gy IR. In panels A and C-E, data are combined from three independent experiments and mean and sem indicated.

**Figure S4.**
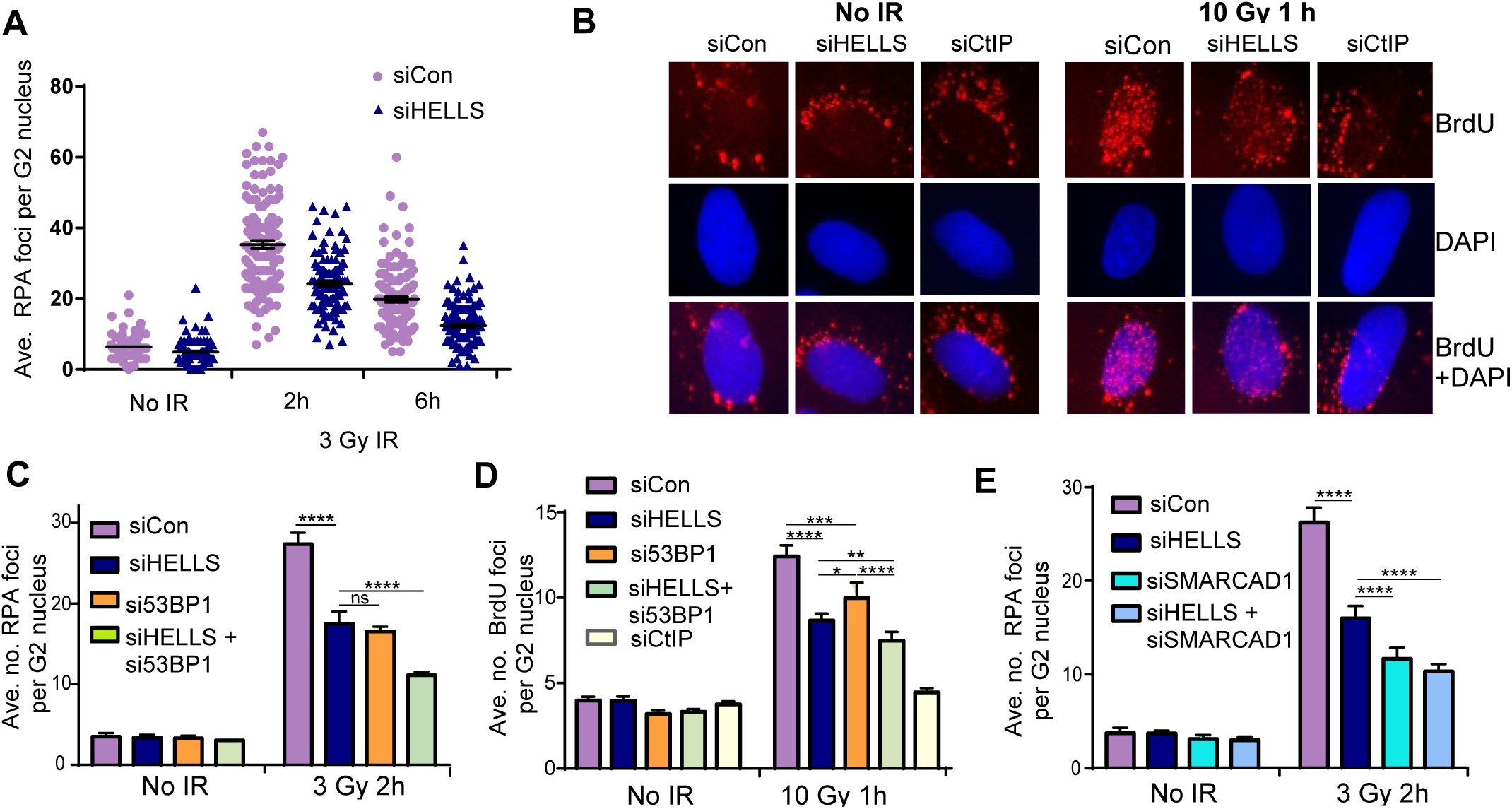
HELLS promotes end resection. **(A)** Individual counts of number of RPA foci per G2 nucleus in 1BR-hTert cells transfected with siCon or siHELLS. **(B)** Representative immunofluorescent images of BrdU foci 1 h after 10Gy IR under non-denaturing conditions in 1BR-hTert cells transfected with the indicated siRNA. **(C)** Quantification of RPA foci per G2 nucleus 2 h after 3Gy, in 1BR-hTert cells transfected with Control, HELLS and 53BP1 siRNAs. **(D)** Quantification of BrdU foci per nucleus 1h after 10 Gy, detected under non-denaturing conditions, in 1BR-hTert cells transfected with Control, HELLS, CtIP and 53BP1 siRNAs. **(E)** Quantification of RPA foci per G2 nucleus 2 h after 3Gy, in 1BR-hTert cells transfected with Control, HELLS and SMARCAD1 siRNAs. **** indicates a p-value of <0.0001, *** indicates p-value of <0.001, ** indicates p-value of <0.01, ** indicates p-value of <0.1 and ns indicates not significant by unpaired two-tailed t-test analysis. For panels A, C, D and E, data shown are the mean +/- sem from three independent experiments.

**Figure S5.**
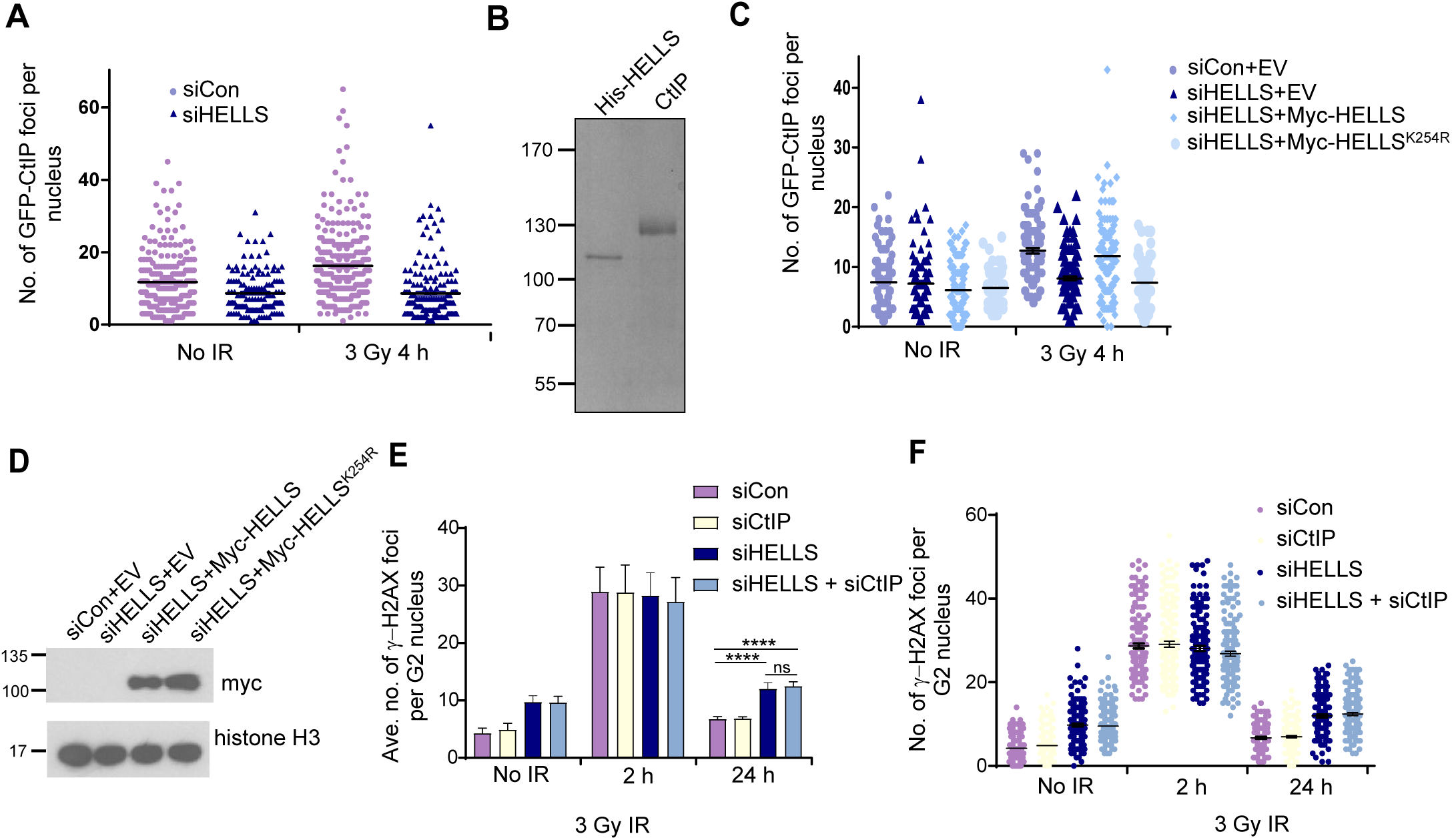
HELLS interacts with CtIP and contributes to its accumulation at IR-induced breaks. **(A)** Individual counts of number of CtIP-GFP per nucleus 4h following 3Gy IR in U2OS cells transfected with CtIP-GFP expressing plasmid and the indicated siRNA. **(B)** Coomassie stained SDS-PAGE gel showing 500 ng of purified recombinant His-HELLS and CtIP proteins. **(C)** Individual counts of the number of IR-induced CtIP-GFP foci per nucleus in U2OS cells transfected with CtIP-GFP expressing plasmid, HELLS siRNA and constructs expressing siRNA-resistant wt or ATPase mutant HELLS. **(D)** Western blot analysis of myc-HELLS wt and myc-HELLS K^254R^ proteins in chromatin fraction of U2OS cells transfected with the indicated siRNA and plasmid. **(E)** Quantification of clearance of γ-H2AX foci in G2 nuclei following 3 Gy IR in 1BR-hTert cells transfected with Control, CtIP and HELLS siRNAs. **(F)** Individual counts of number of γ-H2AX foci per G2 nucleus following 3 Gy IR in 1BR-hTert cells transfected with Control, CtIP and HELLS siRNAs. **** indicates a p-value of <0.0001, and ns indicates not significant by unpaired

